# Shank3 Modulates Sleep and Expression of Circadian Transcription Factors

**DOI:** 10.1101/465799

**Authors:** Ashley M. Ingiosi, Taylor Wintler, Hannah Schoch, Kristan G. Singletary, Dario Righelli, Leandro G. Roser, Davide Risso, Marcos G. Frank, Lucia Peixoto

**Author notes:** To whom correspondence should be addressed: PO Box 1495. Spokane, WA 99210. USA.

## Abstract

Autism Spectrum Disorder (ASD) is the most prevalent neurodevelopmental disorder in the United States and often co-presents with sleep problems. Sleep problems in ASD predict the severity of ASD core diagnostic symptoms and have a considerable impact on the quality of life of caregivers. Little is known, however, about the underlying molecular mechanisms. We investigated the role of *Shank3*, a high confidence ASD gene candidate, in sleep architecture and regulation. We show that mice lacking exon 21 of *Shank3* have problems falling asleep even when sleepy. Using RNA-seq we show that sleep deprivation increases the differences in gene expression between mutants and wild types, downregulating circadian transcription factors *Per3*, *Dec2*, *Hlf*, *Tef*, and *Reverbα*. Shank3 mutants also have trouble regulating wheel-running activity in constant darkness. Overall our study shows that *Shank3* is an important modulator of sleep and clock gene expression.

## Introduction

Autism Spectrum Disorder (ASD) is the most prevalent neurodevelopmental disorder in the United States (diagnosed in 1 in 68 children (Christensen et al., 2016)). The core symptoms of ASD include social and communication deficits, restricted interests, and repetitive behaviors (American Psychiatric Association, 2013). In addition, several studies show that individuals with ASD report a variety of co-morbid conditions including sleep problems. It is estimated that 40 – 80% of the ASD population experience sleep disorders that do not improve with age (Johnson et al., 2009). More specifically, people with ASD have problems falling and staying asleep (Hodge et al., 2014). A recent study showed that sleep problems co-occur with autistic traits in early childhood and increase over time, suggesting that sleep problems are an essential part of ASD (Verhoeff et al., 2018). Indeed, sleep impairments are a strong predictor of the severity of ASD core symptoms as well as aggression and behavioral problems (Cohen et al., 2014; Tudor et al., 2012). Although a great number of studies documented sleep problems in ASD, little is known about the underlying molecular mechanisms.

To better understand the mechanisms underlying sleep problems in ASD, we need animal models that closely recapitulate sleep phenotypes observed in clinical populations. The study of genetic animal models of ASD, in which a genetic abnormality that is known to be associated with ASD is introduced, has provided valuable insight into the molecular mechanisms underlying ASD. These models include Fragile X syndrome, 16p11.2 deletion syndrome, cortical dysplasia-focal epilepsy (CDFE) syndrome and mutations in neuroligins, neurexins and shank genes among others. Sleep research in animal models of ASD is limited and has not yet revealed the underlying mechanisms. Studies using a fly model of Fragile X syndrome reported an increase in sleep which is in contrast to what is observed in the clinical population (Bushey et al., 2009). The opposite phenotype was reported in a Fragile X mouse model, displaying instead an age-dependent reduction in activity during the light phase (i.e the mouse subjective night) (Boone et al., 2018). Mice with a missense mutation in Neuroligin 3 show normal sleep behavior (Liu et al., 2017), but neuroligin 1 knockout mice spend more time asleep (El Helou et al., 2013). Mutant rat models of CDFE syndrome show longer waking periods while the mutant mice show fragmented wakefulness (Thomas et al., 2017). Mice carrying a 16p11.2 deletion syndrome sleep less than wild-type mice, but only males are affected (Angelakos et al., 2017). More importantly, problems of sleep onset, the most prominent feature of sleep problems in ASD patients, have not been evaluated in animal models of ASD.

In this study, we examined sleep in Phelan-McDermid syndrome (PMS) patients with Shank3 mutations and in a mutant mouse with a deletion in Shank3 exon 21 (Shank3^ΔC^). PMS is a syndromic form of ASD characterized by gene deletions affecting the human chromosomal region 22q13.3 (Phelan and McDermid, 2012), particularly the neuronal structural gene *Shank3*. Individuals with PMS have high rates of intellectual disability, difficulties in communication and motor function, and approximately 84% fit the core diagnostic criteria for ASD (Soorya et al., 2013). There is also a high rate of sleep problems in PMS (Bro et al., 2017). Mice with mutations in *Shank3* recapitulate multiple features of both ASD and PMS (Bozdagi et al., 2010; Dhamne et al., 2017; Jaramillo et al., 2017, 2016; Kouser et al., 2013; Peça et al., 2011; Speed et al., 2015), including cognitive impairment, deficits in social behavior, and impaired motor coordination. We show that PMS patients have trouble falling and staying asleep similar to what is observed in the general ASD population. We also show that Shank3^ΔC^ mice sleep less than wild type mice when sleep pressure is high, have reduced sleep intensity (using an accepted electroencephalographic (EEG) metric) under baseline conditions and have delayed sleep onset following sleep deprivation. Our genome-wide gene expression studies show that sleep deprivation sharply increases the differences in gene expression between mutants and wild types, downregulating circadian transcription factors *Per3*, *Dec2*, *Hef*, *Tlf*, and *Reverbα*. We also show that Shank3^ΔC^ mice are unable to sustain wheel-running activity in constant darkness. Overall we show that Shank3 is an important modulator of sleep that may exert its effect through the regulation of circadian transcription factors. Our findings may lead to a deeper understanding of the molecular mechanisms underlying sleep problems in ASD. This may one day lead to the development of successful treatments or interventions for this debilitating comorbidity.

## Results

### Phelan McDermid syndrome patients have problems falling and staying asleep

Recent studies suggest that sleep problems may be present in a substantial number of PMS patients and may be an important factor for caregiver’s well-being (Bro et al., 2017). We obtained genetic and sleep questionnaire data from the Phelan McDermid Foundation International Registry (PMSIR) to estimate the frequency and age of onset of sleep problems in PMS individuals carrying a Shank3 deletion. In parallel, we carried out a meta-analysis of clinical literature to estimate the prevalence of sleep problems in ASD and typically developing populations. Figure 1 shows that PMS patients have trouble falling asleep and experience multiple night awakenings starting at about 5 years of age. Those difficulties translate to reduced time asleep, particularly during adolescence. Although total sleep time seems to improve in adulthood, that improvement seems to be accompanied by an increase in parasomnias. Problems falling and staying asleep persist regardless of age. The frequency of problems falling and staying asleep in PMS patients is similar to what is observed in the general ASD population and much higher than in typically developing individuals (Figure 1 – table supplement 1).

**Figure 1.**
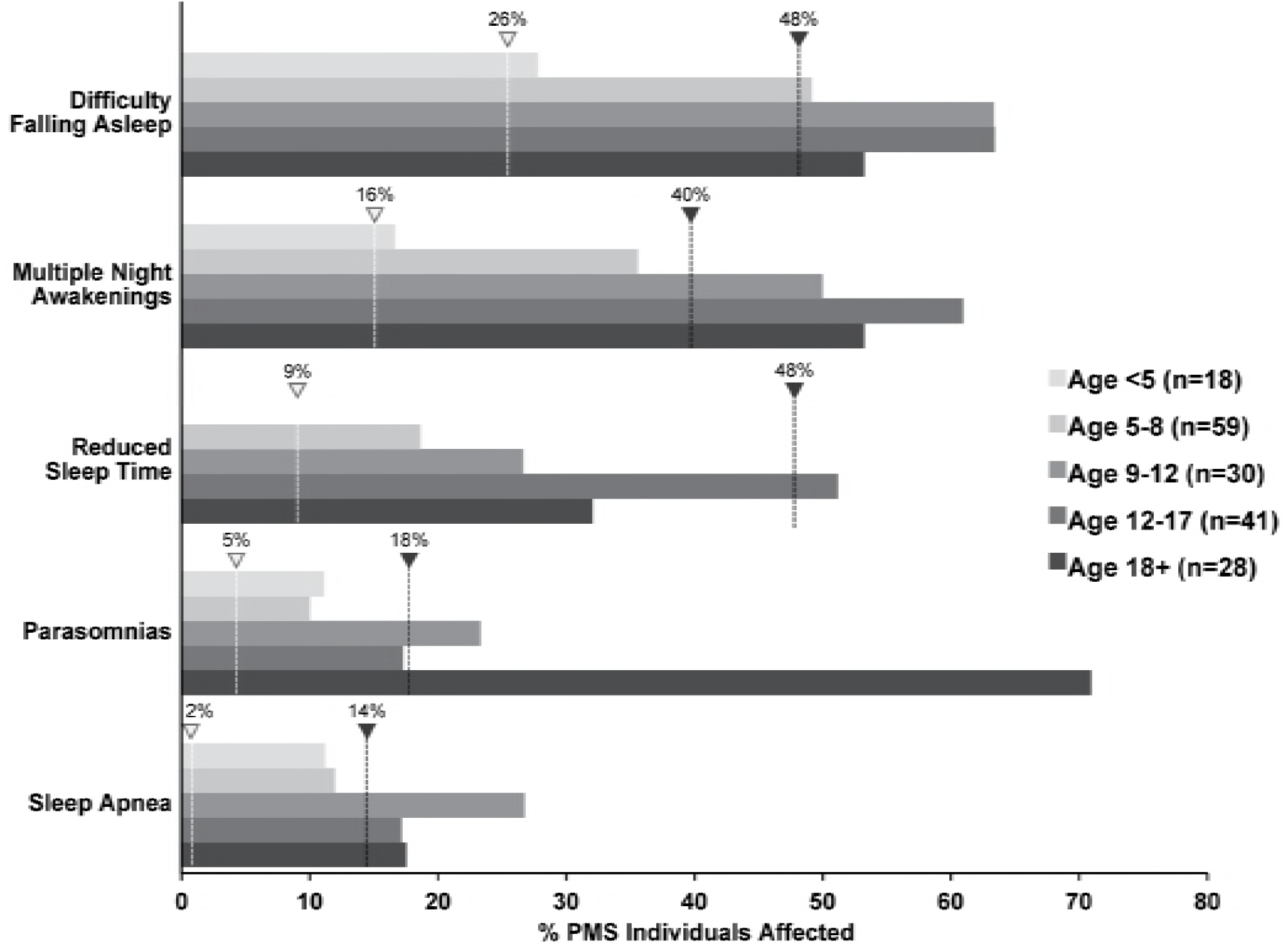
Increased incidence of sleep problems reported in individuals with Phelan-McDermid syndrome (PMS) compared to typically developing (TD) individuals. Dashed line indicates median incidence observed in TD (white marker) and ASD (black marker) populations (values from Figure 1 - table supplement 1).

### Shank3^ΔC^ mice sleep less during the dark phase and show reduced sleep intensity

To determine if Shank3^ΔC^ mice had deficits in spontaneous sleep, undisturbed baseline EEG and electromyographic (EMG) recordings were obtained from wild type (WT) and Shank3^ΔC^ mice. There were no differences between WT and Shank3^ΔC^ mice in time spent in wakefulness or sleep (Figure 2A), bout number (Figure 2 – figure supplement 1A), or bout duration (Figure 2 – figure supplement 1B) in the light period. The two exceptions were that 1) there was a time × genotype effect for REM sleep during hours 1 – 6 and 2) REM bouts were of longer duration in Shank3^ΔC^ mice during hours 7 – 12 (Figure 2 – figure supplement 1B). There were also no differences in time in wakefulness or sleep (Figure 2A) or bout measures (Figure 2 – figure supplement 1A and B) for the first half of the dark period (hours 13 – 18). However during hours 19 – 24, Shank3^ΔC^ mice spent more time in wakefulness and less time in NREM and REM sleep compared to WT mice (Figure 2A). Shank3^ΔC^ mice also had longer waking bouts, shorter NREM and REM bouts, and fewer REM bouts (Figure 2 – figure supplement 1A and B). These data show that Shank3^ΔC^ mice spend more time awake at the end of the dark phase compared to WT mice under baseline conditions.

**Figure 2.**
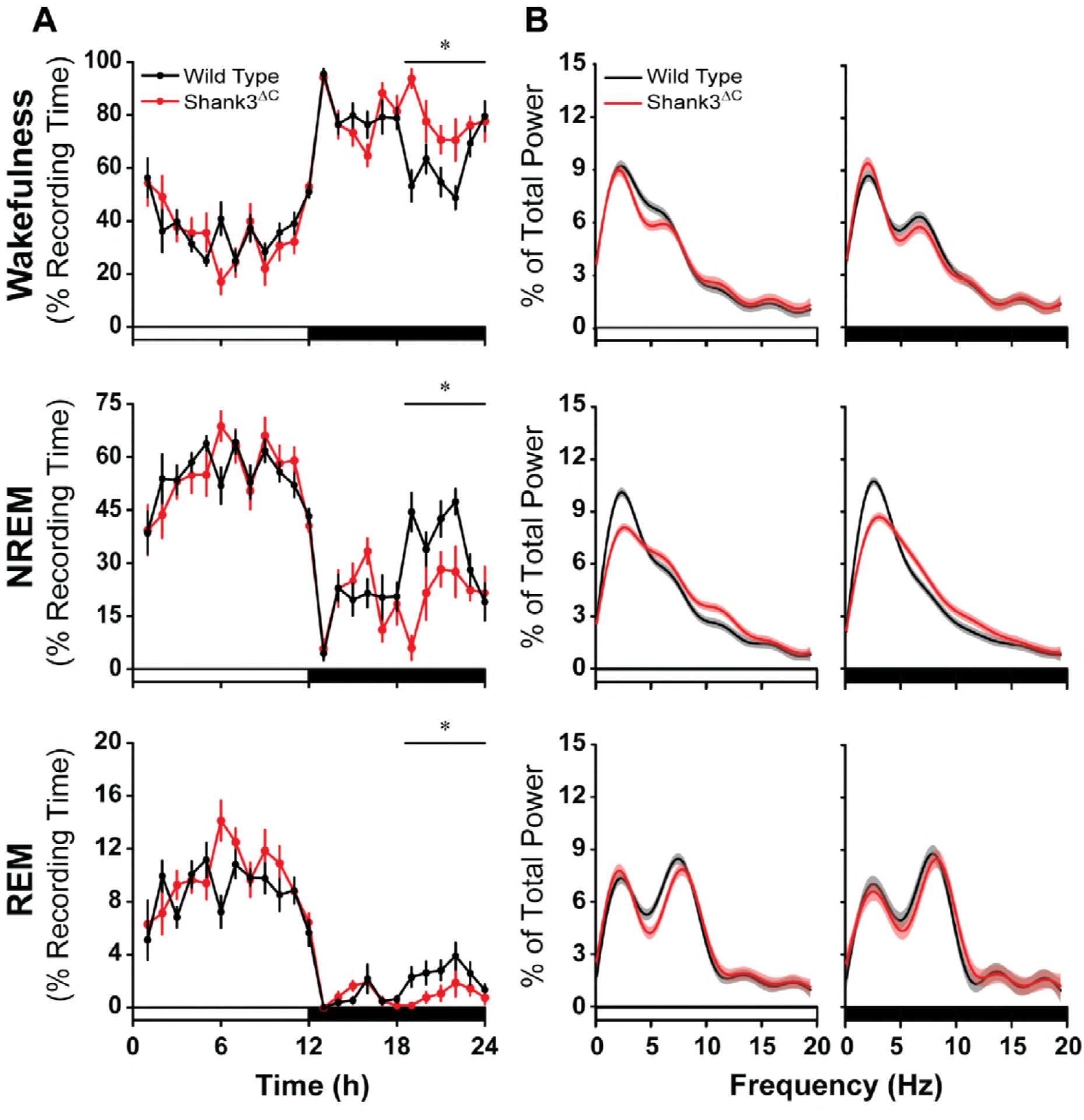
Shank3^ΔC^ mice sleep less during the dark period and show altered EEG spectral power under baseline conditions. The rows represent the arousal states of wakefulness (top), NREM sleep (middle), and REM sleep (bottom). (**A**) Time spent in wakefulness and sleep shown as percentage of recording time per hour. Values are means ± SEM. A significant time × genotype effect was detected for REM sleep in hours 1 – 6 (p = 0.006), but not over the12 hour light phase. (**B**) EEG spectral power normalized as a percentage of the total state-specific EEG power for the light period (left) and dark period (right) fit to smooth curves (solid lines) and expressed with 95% confidence intervals (gray and red shading). The open bars on the x-axis denote the light period and the filled bars denote the dark period of the light:dark cycle. Wild type (n = 10) and Shank3^ΔC^ (n = 10) mice, *p < 0.05.

Fourier analysis of the EEG indicates that Shank3^ΔC^ mice also sleep differently than WT mice (Figure 2B and Figure 2 – table supplement 1). During wakefulness, Shank3^ΔC^ mice showed enhanced power in the alpha band during the light period as well as enhanced delta power and blunted theta power during the dark period. During NREM sleep, delta power was blunted and theta and alpha bands were enhanced for Shank3^ΔC^ mice during the light and dark periods. During REM sleep, the alpha band was elevated in Shank3^ΔC^ mice compared to WT during the light period. Because NREM delta power is a measure of sleep intensity or depth (Achermann and Borbely, 2017), these data suggest that Shank3^ΔC^ mice sleep less deeply under baseline conditions.

### Shank3^ΔC^ mice have abnormal homeostatic responses to sleep deprivation

To investigate sleep homeostasis in Shank3^ΔC^ we sleep deprived mutant and WT mice for 5 hours via gentle handling starting at light onset and sacrificed immediately post-SD. Both WT and Shank3^ΔC^ mice showed homeostatic responses to SD (Figure 3 – figure supplement 1) with reduced wakefulness and increased time spent in NREM and REM sleep during the recovery phase. This response was delayed until the dark period in Shank3^ΔC^ mice (Figure 3 – figure supplement 1B). Sleep was more consolidated after SD for both WT and Shank3^ΔC^ mice (Figure 3 – figure supplement 2A – B), but Shank3^ΔC^ mice had fewer NREM bouts/h for hours 13 – 18. In the last 6 h of the dark period, Shank3^ΔC^ mice spent more time awake and less time in NREM and REM sleep post-SD (Figure 3 – figure supplement 1C) which is consistent with baseline data (Figure 2A). Overall, Shank3^ΔC^ mice showed homeostatic increases in sleep and consolidation of arousal states during recovery sleep, but this response was delayed in Shank3^ΔC^ mice compared to WT animals (Figure 3 – figure supplement 1 – 2). Changes in NREM EEG delta power were similar between WT and Shank3^ΔC^ mice, indicating that mutant mice accumulate sleep pressure similarly to WT mice (Figure 3A). However, Shank3^ΔC^ mice showed a transient enhancement of power in the higher frequencies (3.9 – 19.5 Hz; Figure 3B) compared to WT mice in the first 2 h post-SD (hours 6 – 7) that resolved to baseline and WT values during hours 11 – 12 (Figure 3C). Overall, these data, as well as depressed NREM baseline delta power, suggest the presence of abnormal sleep homeostasis in Shank3^ΔC^ mice.

**Figure 3.**
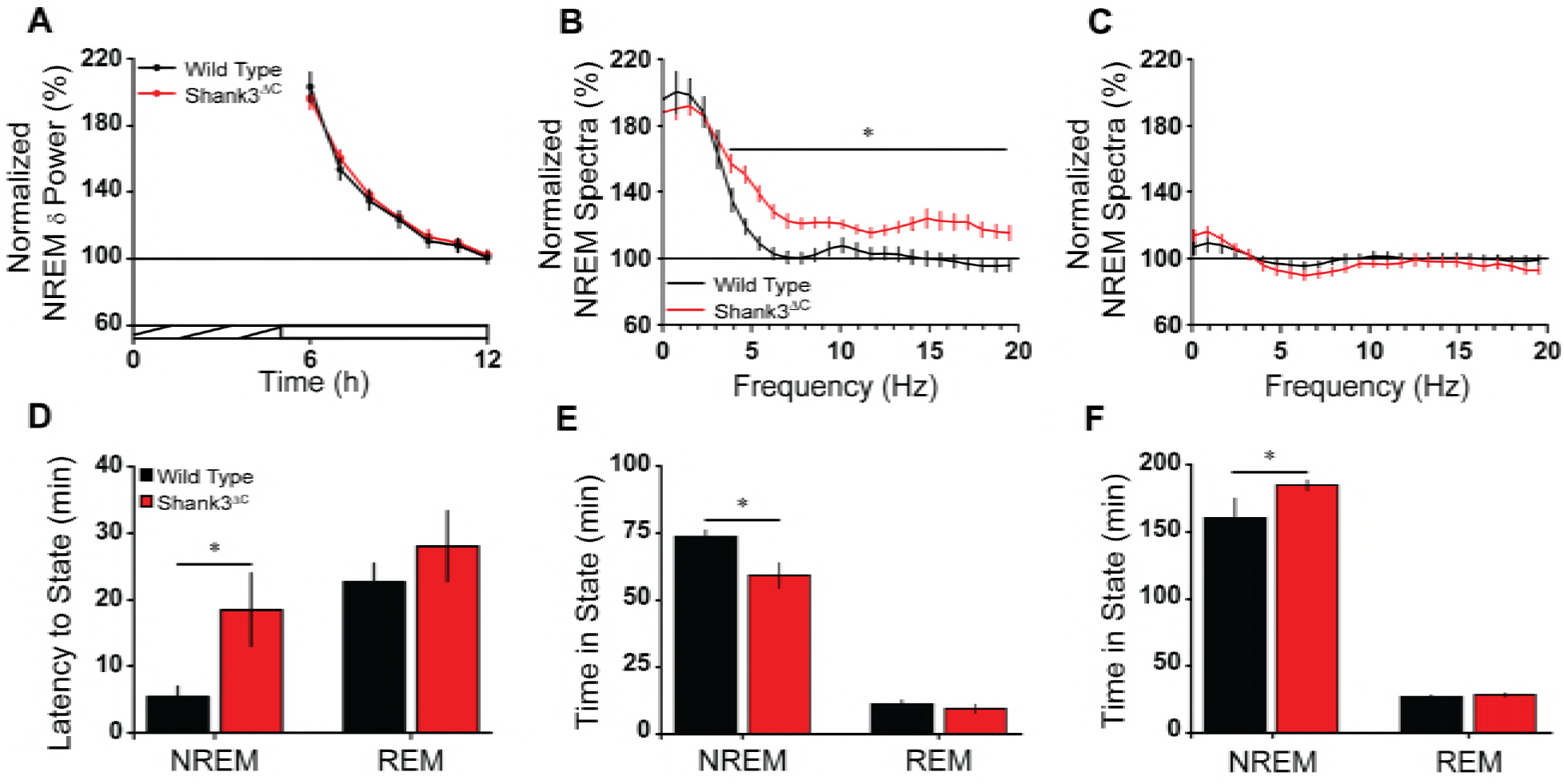
Shank3^ΔC^ mice take longer to fall asleep and sleep less after sleep deprivation. (**A**) Normalized NREM sleep delta power as measured by power in the EEG delta (δ; 0.5 – 4 Hz) frequency band relative to baseline. The cross-hatched bar of the x-axis denotes the 5 h sleep deprivation period and the open bar denotes the remaining light period. (**B**) Normalized NREM spectra for the first 2 h post-sleep deprivation (hours 6 – 7; significant from 3.9 – 19.5 Hz). (**C**) Normalized NREM spectra for the remaining 2 h of light period post-sleep deprivation (hours 11 – 12). (**D**) Latency to enter NREM sleep and REM sleep after sleep deprivation. (**E**) Time in NREM sleep and REM sleep for the first 2 h post-sleep deprivation (hours 6 – 7). (**F**) Time in NREM sleep and REM sleep for the remaining 5 h of light period post-sleep deprivation (hours 8 – 12). Values are means ± SEM for wild type (n = 10) and Shank3^ΔC^ (n = 10) mice, *p < 0.05.

Remarkably, Shank3^ΔC^ mice took longer to enter NREM sleep post-SD compared to WT (Figure 3D). As a consequence, Shank3^ΔC^ mice spent less time in NREM sleep compared to WT during the first 2 h of recovery sleep (Figure 3E). During the subsequent 5 h of the light period (hours 8 – 12), Shank3^ΔC^ mice spent more time in NREM sleep compared to WT (Figure 3F). Overall our data show that Shank3^ΔC^ mice have difficulties falling asleep despite heightened sleep pressure.

### Shank3^ΔC^ mice show downregulation of circadian transcription factors in response to sleep deprivation

We conducted a genome-wide gene expression study to investigate the molecular basis for the Shank3^ΔC^ mouse phenotype Shank3^ΔC^ and WT adult male mice were subjected to 5 hours of SD starting at light onset. Mice from both genotypes were sacrificed at the same time of day to determine differences in gene-expression under home cage conditions (HC). Prefrontal cortex was collected for all animals and subjected to RNA sequencing (RNA-seq, n = 5 per group). As expected, sleep deprivation is the greatest source of variation in the data, followed by the genotype effect (Figure 4A). Furthermore, the difference between genotypes is increased after sleep deprivation (Figure 4A), which greatly increases the number of differentially expressed genes between Shank3^ΔC^ and WT mice, starting at 67 genes (HC) and doubling to 136 genes (SD) (Figure 4B, Figure 4 – table supplement 1; FDR < 0.5) Most of the differences in gene expression following SD are not present in HC conditions.

**Figure 4.**
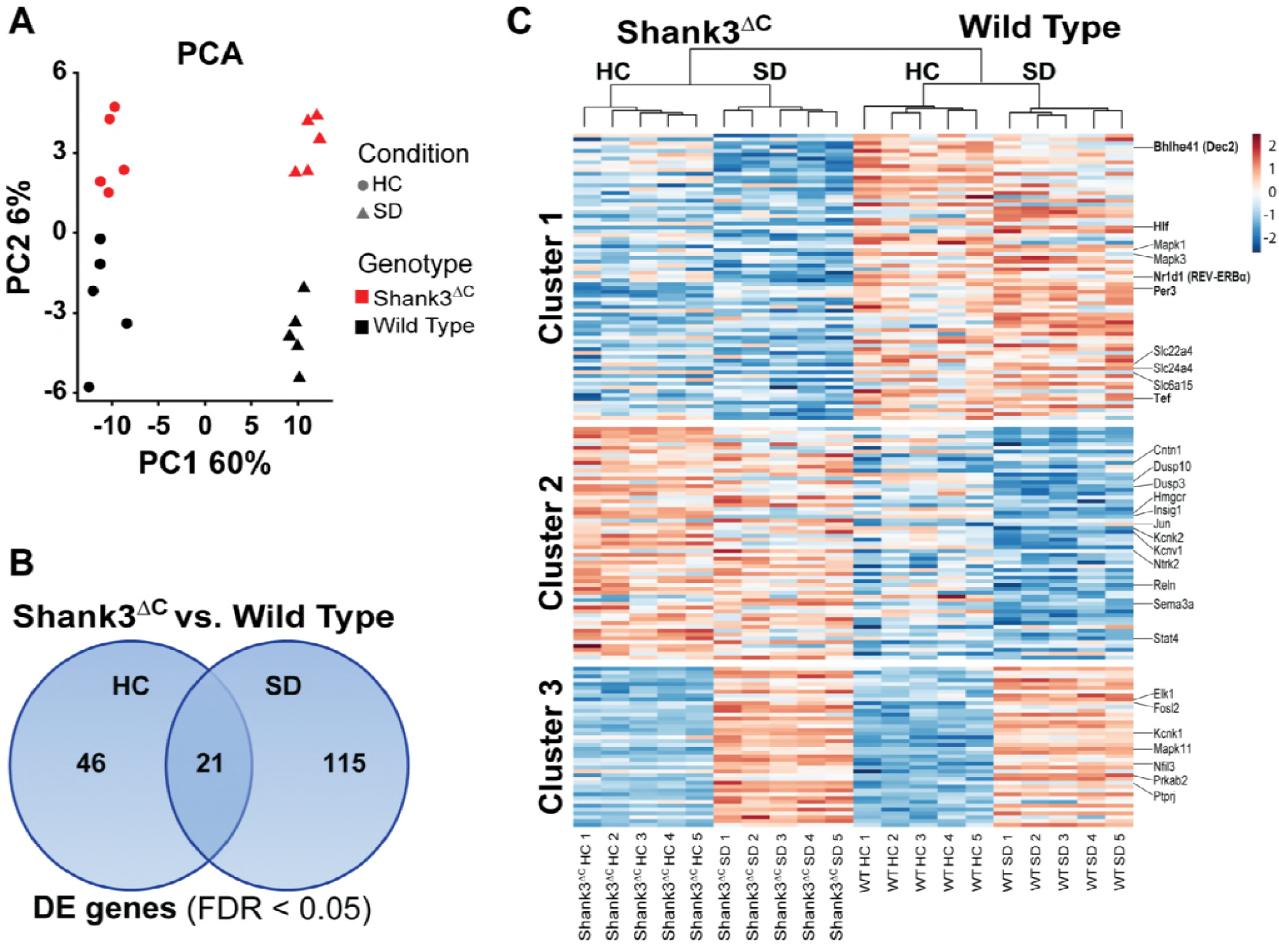
Sleep deprivation induces a two-fold difference in gene expression between Shank3^ΔC^ and wild type mice. RNA-seq study of gene expression from prefrontal cortex obtained from adult male Shank3^ΔC^ and wild type mice, either under control home cage conditions (HC) or following 5 hours of sleep deprivation (SD). N=5 mice per group. (**A**) Principal component analysis of normalized RNA-seq data shows that sleep deprivation is the main source of variance in the data (first principal component, PC1) and genotype is the second (second principal component, PC2). Percent variance explained by each PC is shown in each axis. (**B**) Venn diagram showing the number of genes differentially expressed at FDR < 0.05 between Shank3^ΔC^ and wild type mice in either control HC conditions or after SD. (**C**) Heat map of average scaled gene expression for all genes in (**B**). K-means clustering defined 3 clusters based on differences in gene expression across all comparisons. Genes belonging to the MAPK pathway and involved in circadian rhythms (see Table 1) are highlighted on the right.

Clustering of gene expression patterns for all genes differentially expressed between Shank3^ΔC^ and WT mice reveals 3 groups of genes (Figure 4C). Cluster 1 contains genes that are downregulated in mutants versus WT mice under HC conditions. Genes in this cluster are also downregulated in response to SD in Shank3^ΔC^ mice but are generally unaffected by SD in WT mice. SD seems to exacerbate the difference between genotypes, which explains why differential expression of some of the genes in cluster 1 only reach statistical significance after SD. This cluster contains the majority of the genes that are differentially expressed between genotypes. Cluster 2 contains genes that are upregulated in mutants versus WT mice under HC conditions that are also downregulated by SD in WT mice. Cluster 3 contains genes that are normally upregulated by SD. For both clusters 2 and 3, the Shank3^ΔC^ mutation seems to dampen the response to SD.

To better understand the impact of the gene expression at the pathway level, we carried out functional annotation of transcripts differentially expressed between Shank3^ΔC^ and WT mice (Table 1, Table 1 – table supplement 1). Our results revealed that MAPK/GnRH signaling and circadian rhythm-associated transcripts are downregulated in Shank3^ΔC^ mice, and that sleep deprivation exacerbates that difference. Circadian transcription factors are particularly affected. For example, while expression of *Per3*, *Hlf* and *Tef* is already different under homecage conditions, SD leads to a genotype specific difference in expression of *Nr1d1* (*Reverbα*) and *Bhlhe41* (*Dec2*). All of the above mentioned circadian transcription factors belong to cluster 1 on our heat map (Figure 4C) showing downregulation in response to SD only in Shank3^ΔC^ mice.

**Table 1.**
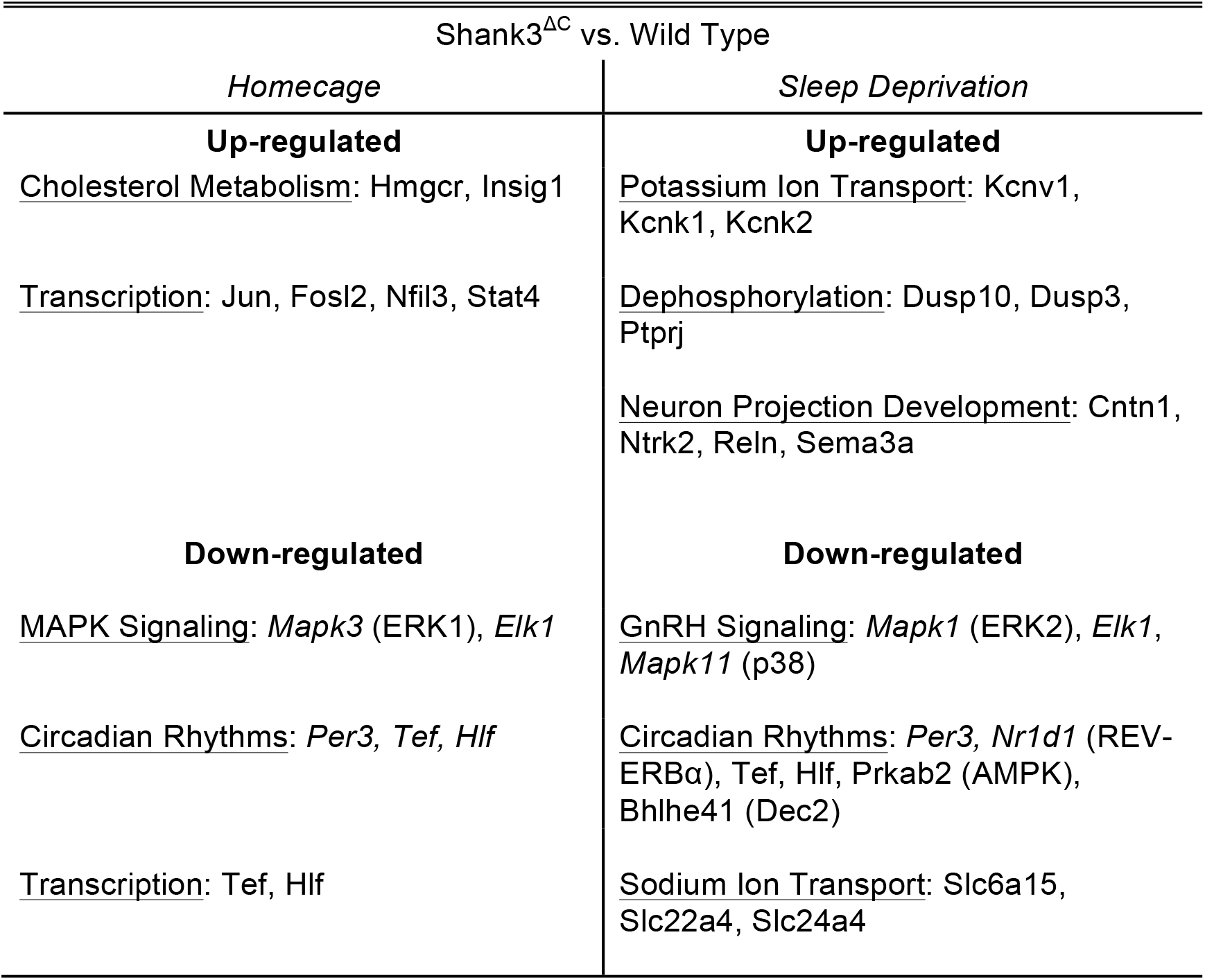
Results of functional bioinformatics analysis of the genes differentially expressed between Shank3^ΔC^ relative to wild type mice. Functional annotation and clustering analysis was performed using DAVID (https://david.ncifcrf.gov) and functional information was obtained from the following databases: GO (Biological process and Molecular function), KEGG pathways, Uniprot keywords. Enrichment was performed relative to all transcripts expressed in the mouse prefrontal cortex as defined by our RNAseq data. Enriched functional terms were clustered at low stringency, to obtain clusters with enrichment score > 1.2 (corresponding to an average p-value > 0.05). See Table 1 – table supplement 1 for details.

### Wheel-running activity in Shank3^ΔC^ mice is impaired in constant darkness

The results of our RNA-seq analysis revealed a general downregulation of circadian transcription factors in the mutants. These findings suggested that Shank3^ΔC^ mice might also have abnormal circadian rhythms. To test this we measured wheel-running activity in Shank3^ΔC^ and WT mice. Wheel running data were collected for 2 weeks under LD (LD weeks 2 – 3; Figure 6 and Figure 6 – figure supplement 1). Alpha, the length of activity period, was calculated as the time between activity onset and offset. Period length and alpha length did not differ between Shank3^ΔC^ and WT mice (Table 2). However, Shank3^ΔC^ mice ran fewer revolutions/day compared to WT (Table 2). Mice were then released into constant darkness (DD) for 3 weeks. There was no difference between Shank3^ΔC^ and WT mice for period length during the DD period (Table 2). Shank3^ΔC^ mice showed significant but small decreases in alpha length compared to WT during DD weeks 1 and 2 (Table 2). Shank3^ΔC^ mice also continued to run less than WT mice for the entire DD period (Table 2). Furthermore, there was a significant genotype × time interaction effect for wheel running activity from LD week 3 – DD week 3 (Table 2, Table 2 - table supplement 1). Some Shank3^ΔC^ mice greatly reduced their running at certain periods of the DD period, which is reflected in variance for period, alpha, and activity measures (Figure 5, Figure 5 – figure supplement 1, and Table 2). Daily checks were made to verify that each running wheel was transmitting data and that no light was detected in the room during constant darkness. Overall, these data indicate that constant darkness impairs wheel-running activity of Shank3^ΔC^ mice.

**Table 2.** Shank3^∆C^ mice show reduced activity during constant darkness. Summary of wheel running behavior measured for each week of 12:12 h light dark cycle (LD, 559 ± 4: 0 ± 0 Lux) or constant darkness (DD, 0 ± 0 Lux) for both wild type and Shank3^∆C^ mice. Comparisons between genotypes (WT vs. Shank3^∆C^) were done with paired or unpaired Student’s t-tests as appropriate for period, alpha, and running wheel activity. A linear mixed-effects model was used to estimate the contributions of genotype and the interaction between genotype and time for period, alpha, and wheel running activity for LD week 3 through DD week 3. There is a significant interaction effect for genotype × time for activity (p < 0.0001, ANOVA; indicated by vertical bar) but no significant effect for genotype × period (p = 0.15, ANOVA) or genotype × alpha (p = 0.88, ANOVA). See Table 2 – table supplement 1 for details. Values are means ± SEM for wild type (n = 8) and Shank3^∆C^ (n = 7) mice, *p < 0.05.

|  | *Wild Type* | *Shank3<sup>ΔC</sup>* |
| --- | --- | --- |
| *Period (h)* |  |  |
| LD Week 2 | 24.0 ± 0.0 | 24.0 ± 0.0 |
| LD Week 3 | 24.0 ± 0.0 | 23.9 ± 0.1 |
| DD Week 1 | 23.9 ± 0.1 | 23.7 ± 0.1 |
| DD Week 2 | 23.7 ± 0.1 | 20.3 ± 3.4 |
| DD Week 3 | 23.5 ± 0.2 | 25.8 ± 1.5 |
| *Alpha (h)* |  |  |
| LD Week 2 | 10.0 ± 0.5 | 10.3 ± 0.5 |
| LD Week 3 | 10.2 ± 0.3 | 10.8 ± 0.5 |
| DD Week 1 | 10.5 ± 0.2 | **9.0 ± 0.4\*** |
| DD Week 2 | 10.4 ± 0.4 | **8.2 ± 0.7\*** |
| DD Week 3 | 9.0 ± 1.0 | 9.9 ± 1.0 |
| *Activity (rev/day)* |  |  |
| LD Week 2 | 33985.2 ± 565.9 | **23258.2 ± 861.2\*** |
| LD Week 3 | 40306.7 ± 1257.1 | **25365.1 ± 964.0\*** |
| DD Week 1 | 37942.6 ± 795.4 | **23027.7 ± 669.6\*** |
| DD Week 2 | 38250.7 ± 431.1 | **16820.7 ± 1302.5\*** |
| DD Week 3 | 38586.3 ± 231.1 | **6961.5 ± 383.9\*** |
| **p < 0.0001** |  |  |

**Figure 5.**
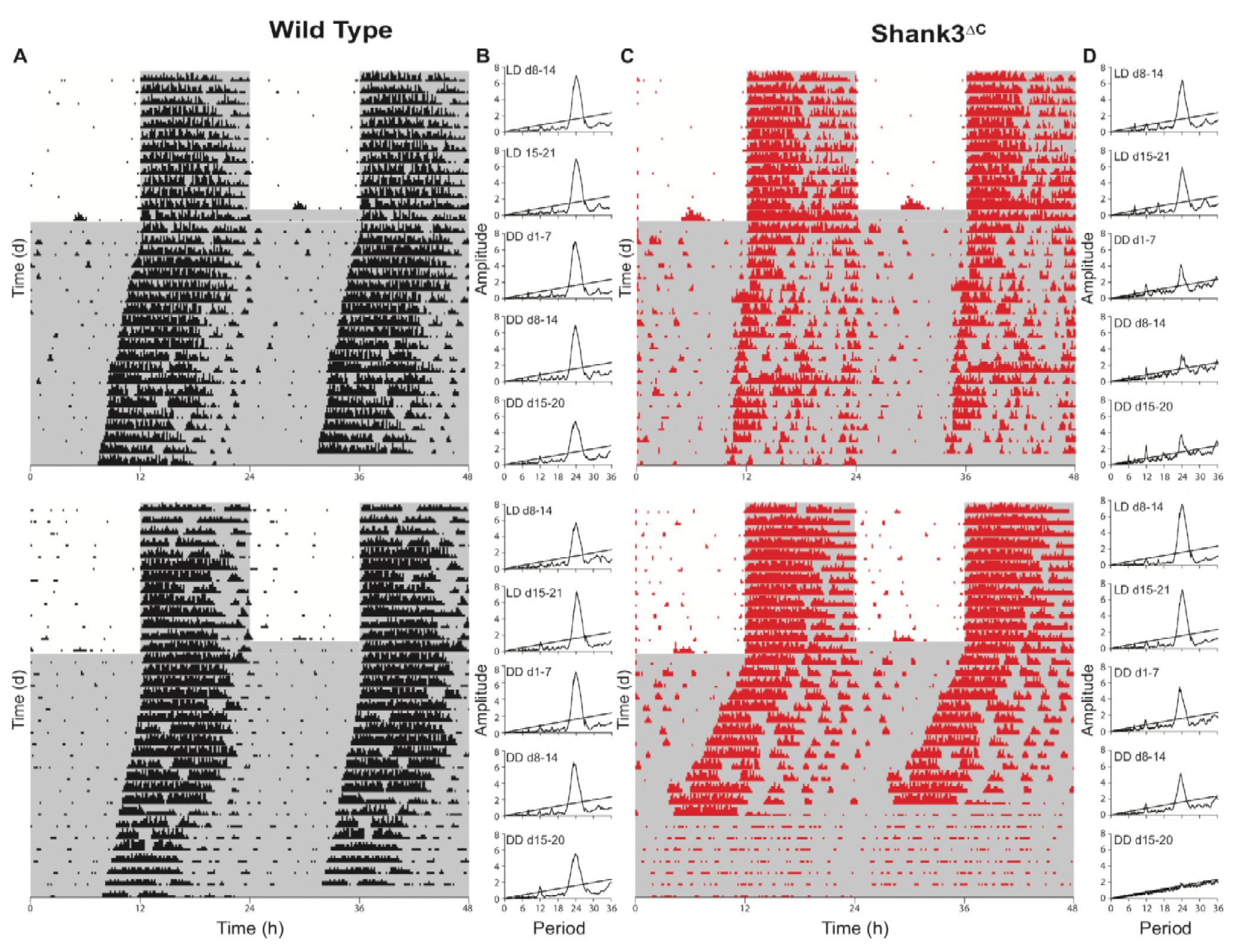
Shank3^ΔC^ mice show disruption of running wheel activity in constant darkness. Representative actograms and periodograms for two wild type and two Shank3^ΔC^ mice. Mice were entrained to a 12:12 h light:dark cycle (LD, 559 ± 4: 0 ± 0 lux) for two weeks prior to 3 weeks constant darkness (DD, 0 ± 0 lux). Gray shading is representative of the dark period. (**A**) Actograms for two wild type mice. (**B**) Corresponding periodograms for wild type mice. (**C**) Actograms for two Shank3^ΔC^ mice. (**D**) Corresponding periodograms for Shank3^ΔC^ mice. **Table 1**.

## Discussion

The present study is the first to establish a role for *Shank3* in mammalian sleep. We show that both PMS patients and *Shank3* mutant mice have trouble falling asleep (Figures 1, 2, and 3). This phenotype is widely observed in the ASD population and until now had not been replicated in animal models. Shank3^∆C^ mice have problems falling asleep when sleep pressure is high, such as the end of the dark period (Figure 2A) or following sleep deprivation (Figure 3D – E). They also sleep less deeply (Figure 2B). However, they show no gross abnormalities in sleep homeostasis or circadian rhythms, (Figure 3A, Figure 5). This suggests that the primary deficit is in sleep onset.

Our molecular studies show that sleep deprivation increases differences in gene expression between Shank3^∆C^ and WT mice (Figure 4). These differences point to the downregulation of circadian transcription factors and genes involved in the MAPK/GnRH pathways in the mutants (Table 1). Circadian transcription factors affected include *Per3*, *Hlf*, *Tef*, *Reverbα* and *Dec2*. We therefore investigated the effects of our *Shank3* mutation in circadian rhythms. Shank3^∆C^ mice do not show a disruption in circadian rhythmicity but a large reduction in wheel-running activity in response to constant darkness (Figure 5, Table 2). This suggests that the mutant sleep phenotype involves clock gene function outside of their central time-keeping role. Interestingly mutations in some of the circadian transcription factors we identified, *Per3* and *Dec2*, indeed lead to deficits in sleep regulation (Archer et al., 2018; He et al., 2009; Hirano et al., 2018). Our results support a role of clock genes in influencing sleep outside of their roles in generating circadian rhythms (Franken, 2013).

An interesting question is how can deletion of exon 21 of Shank3 lead to dysregulation of transcriptional and signaling pathways linked to sleep and sleep loss? Exon 21 of Shank3 encodes the homer and cortactin interaction domains of the protein. Homer interacts with metabotropic glutamate receptors (mGluRs) and Shank3/homer complexes anchor mGluRs to the synapse. Shank3^∆C^ mice show a marked reduction of the major isoforms of Shank3 as well as an increase of mGluRs at the synapse (Kouser et al., 2013). mGluR signaling activates the MAPK/ERK pathway, a key regulator of activity-dependent transcription and synaptic plasticity in mature neurons (Thomas and Huganir, 2004). So the role of SHANK3 at the synaptic membrane explains the observed regulation of MAPK pathway genes. However, circadian transcription factors act within the nucleus. Interestingly, SHANK3 is known to undergo synaptic-nuclear shuttling in response to neuronal activity and interact with nuclear ribonucleoproteins and components of the RNA Pol II mediator complex (Grabrucker et al., 2014). Deletion of the c-terminus leads to nuclear accumulation of SHANK3 and alterations in gene expression (Cochoy et al., 2015; Grabrucker et al., 2014). Thus, mutations in exon 21 of *Shank3* could lead to deficits in transcriptional regulation in response to sleep deprivation through direct regulation of transcription in the nucleus. Yeast-two hybrid data shows that SHANK3 can directly bind the circadian transcription factors REVERBα (NR1D1) and DEC1 (BHLHE40), a close paralog of DEC2 (Sakai et al., 2011). Therefore, we propose that SHANK3 acts as a scaffolding protein in the nucleus that helps assemble protein complexes that are key for the regulation of sleep onset via circadian transcription factors.

## Materials and methods

### Sleep questionnaire study

Sleep questionnaire data for PMS patients was obtained through a data access agreement with the PMSIR. The PMSIR contains demographic, clinical and genetic data on PMS patients. Parents and caregivers are able to create a profile on behalf of the patient, enter demographic data, complete questionnaires on symptoms and development, and upload files such as genetic test reports and other medical records. Trained genetic counselors curate all genetic data to ensure each patient has as complete and accurate of a “genetic profile” as possible. Researchers may apply for data exports containing de-identified clinical data, developmental data and genetic data. Once approved, the appropriate search is performed in the database and the results are provided to the researcher. Data in this study corresponds to a database export performed on 12/1/2016 containing the results of a sleep questionnaire completed by caregivers, as well as biographic and genetic information of PMS individuals. Problems with sleep onset are defined as more than an hour to fall asleep. Multiple night awakenings are defined as more than two. Reduced sleep time is defined as less than 6 hours per night. Parasomnias are defined as abnormal movements, behaviors, emotions, perceptions, and dreams that occur while falling asleep or sleeping. Presence of sleep apnea is defined as having received a diagnosis. Only individuals with a genetic counselor confirmed deletion of the *Shank3* gene were included in the analysis. The final dataset included 176 individuals, 78 males and 98 females, age range 1 – 39 years.

### Animals

Heterozygous Shank3^+/∆C^ mice were obtained from Dr. Paul Worley at Johns Hopkins University. Shank3^+/∆C^ breeding pairs were established to obtain wild type (WT) and Shank3^∆C^ littermates. Mice were housed in standard cages at 24 ± 1°C on a 12:12 h light:dark cycle with food and water *ad libitum*. All experimental procedures were approved by the Institutional Animal Care and Use Committee of Washington State University and conducted in accordance with National Research Council guidelines and regulations for experiments in live animals.

### Assessment of sleep-wake behavior

#### Surgical procedures

Adult male mice (12-weeks-old) were stereotaxically implanted with electroencephalographic (EEG) and electromyographic (EMG) electrodes under isoflurane anesthesia according to previously published methods (Frank et al., 2002). Briefly, four stainless steel screws (BC-002MPU188, Bellcan International Corp, Hialeah, FL) were placed contralaterally over frontal (2) and parietal (2) cortices, and 2 EMG electrodes were inserted in the nuchal muscles. Mice were allowed 5 days of recovery from surgery prior to habituation to the recording environment.

#### Experimental design

Six days after surgery, wild type (WT; n = 10) and Shank3^∆C^ (n = 10) littermates were connected to a lightweight, flexible tether and allowed 5 days to habituate to the tether and recording environment. After habituation, mice underwent 24 h undisturbed baseline EEG and EMG recording starting at light onset. The next day, mice were sleep deprived for 5 h via gentle handling beginning at light onset according to previously published methods (Halassa et al., 2009; Vecsey et al., 2012; Ingiosi et al., 2015). Mice were then allowed 19 h undisturbed recovery sleep.

#### EEG / EMG data acquisition and analysis

EEG and EMG data were collected with Grass 7 polygraph hardware (Natus Medical Incorporated, Pleasanton, CA) via a light-weight, counterbalanced cable, amplified, and digitized at 200 Hz using VitalRecorder acquisition software (SleepSign for Animal, Kissei Comtec Co., LTD, Nagano, Japan). EEG and EMG data were high- and low-pass filtered at 0.3 and 100 Hz and 10 and 100 Hz, respectively.

Wakefulness and sleep states were determined by visual inspection of the EEG waveform, EMG activity, and fast Fourier transform (FFT) analysis by an experimenter blinded to the genotype. Data were scored as wakefulness, NREM sleep, or REM sleep with 4-s resolution using SleepSign for Animal as previously described (Halassa et al., 2009).

Statistical analyses were performed using SPSS for Windows (IBM Corporation, Armonk, NY) and R statistical language. Data are presented as means ± standard error of the mean (SEM). A general linear model for repeated measures using time (hours) as the repeated measure and genotype (WT vs. Shank3^∆C^) or manipulation (undisturbed baseline vs. 5 h sleep deprivation) as the between subjects factor was used when multiple measurements were made over time (i.e. time in stage, hourly NREM delta power). Comparisons between genotypes (WT vs. Shank3^∆C^) or manipulations (undisturbed baseline vs. 5 h sleep deprivation) were performed with paired or unpaired Student’s t-test as appropriate for bout analyses, spectral bands, latency to state, and total time in stage. Latency to NREM or REM sleep after sleep deprivation was defined as time elapsed from release to recovery sleep to the first bout of NREM sleep (bout ≥ 28 s or 7 consecutive epochs) or REM sleep (bout ≥ 16 s or 4 consecutive epochs).

The EEG was subjected to fast Fourier transform analysis to produce power spectra between 0 – 20 Hz with 0.781 Hz resolution. Delta (δ) was defined as 0.5 – 4 Hz, theta (θ) as 5 – 9 Hz, and alpha (α) as 10 – 15 Hz. For comparison of light and dark period spectral data, each spectral bin was expressed as a percentage of the total state-specific EEG power (0 – 20 Hz) of the baseline 12 h light period and 12 h dark period, respectively (Baracchi and Opp, 2008). The relation between EEG spectral power and frequency under baseline conditions was fit to smooth curves and 95% confidence intervals for the respective light and dark periods. The analysis was performed via Generalized Additive Models (GAM) (T.J. Hastie and R.J. Tibshirani, 1990) using the R package mgcf (v. 3.5.0). For hourly NREM delta power analysis, spectral values within the delta band for each hour were normalized to the average NREM delta band value of the last 4 h of the baseline light period (zeitgeber time (ZT) 8 – 11) and expressed as a percentage (Franken et al., 2001). Changes in individual NREM EEG spectral bins (0 – 20 Hz) were normalized to corresponding baseline spectral bins obtained from mean values from the last 4 h of the baseline light period (ZT8 – 11). EEG epochs containing artifacts were excluded from spectral analyses.

### Genome-wide gene expression

#### RNA isolation

Adult male (8 – 10-week-old) Shank3^∆C^ mice and WT littermates were divided into 2 groups: homecage controls (WT n = 5; Shank3^∆C^ n =5) and sleep deprived (WT n = 5; Shank3^∆C^ n =5). All mice were individually housed. Homecage control mice were left undisturbed and sacrificed 5 h after light onset (ZT4). Mice in the sleep deprived group were sleep deprived for 5 h via gentle handling starting at light onset and then sacrificed upon completion of sleep deprivation (ZT4) without allowing for any recovery sleep. Mice were sacrificed by live cervical dislocation (alternating between homecage controls and sleep deprived mice), decapitated, and prefrontal cortex was swiftly dissected on a cold block. Tissue was flash frozen in liquid nitrogen at and stored at −80°C until processing (Poplawski et al., 2016) (Tudor et al., 2016). This protocol was repeated over a 5 day period, and all tissue was collected within the first 15 min of the hour. Tissue was later homogenized in Qiazol buffer (Qiagen, Hilden, Germany) using a TissueLyser (Qiagen) and all RNA was extracted using the Qiagen RNAeasy kit (Qiagen) on the same day.

#### RNA-seq library preparation and sequencing

The integrity of total RNA was assessed using Fragment Analyzer (Advanced Analytical Technologies, Inc., Ankeny, IA) with the High Sensitivity RNA Analysis Kit (Advanced Analytical Technologies, Inc.). RNA Quality Numbers (RQNs) from 1 to 10 were assigned to each sample to indicate its integrity or quality. “10” stands for a perfect RNA sample without any degradation, whereas “1” marks a completely degraded sample. RNA samples with RQNs ranging from 8 to 10 were used for RNA library preparation with the TruSeq Stranded mRNA Library Prep Kit (Illumina, San Diego, CA). Briefly, mRNA was isolated from 2.5 µg of total RNA using poly-T oligo attached to magnetic beads and then subjected to fragmentation, followed by cDNA synthesis, dA-tailing, adaptor ligation and PCR enrichment. The sizes of RNA libraries were assessed by Fragment Analyzer with the High Sensitivity NGS Fragment Analysis Kit (Advanced Analytical Technologies, Inc.). The concentrations of RNA libraries were measured by StepOnePlus Real-Time PCR System (ThermoFisher Scientific, San Jose, CA) with the KAPA Library Quantification Kit (Kapabiosystems, Wilmington, MA). The libraries were diluted to 2 nM with Tris buffer (10 mM Tris-HCl, pH8.5) and denatured with 0.1 N NaOH. Eighteen pM libraries were clustered in a high-output flow cell using HiSeq Cluster Kit v4 on a cBot (Illumina). After cluster generation, the flow cell was loaded onto HiSeq 2500 for sequencing using HiSeq SBS kit v4 (Illumina). DNA was sequenced from both ends (paired-end) with a read length of 100 bp.

#### RNA-seq data analysis

HiSeq Control Software version 2.2.68 was used for base calling. The raw bcl files were converted to fastq files using the software program bcl2fastq2.17.1.14. Adaptors were trimmed from the fastq files during the conversion. The average sequencing depth for all samples was 52 million read pairs. Sequenced reads were mapped with GSNAP (parameters = -N 1 -m 7 -A sam —nofails) to mm10. On average 84% of the sequenced reads mapped uniquely to the mm10 genome. All statistical analyses were performed using open source software freely available through the R/Bioconductor project (Gentleman et al., 2004). The count matrix was obtained using featureCounts (parameters = isPairedEnd = TRUE, requireBothEndsMapped = TRUE, annot.inbuilt = "mm10") from the package Rsubread (v. 1.26.1) under Bioconductor (version 3.6) with R (v. 3.4.1) using mm10 built-in annotations (Mus_musculus.GRCm38.90) with ENTREZ ID as row names. Gene counts were filtered using a proportion test (counts per million cutoff of 1), as implemented in the NOISeq package (v.2.22.1) (Tarazona et al., 2015). Removal of unwanted variation (RUV) normalization was performed using RUVSeq (v. 1.12.0) (Risso et al., 2014) after the data was normalized by Trimmed Mean of M-values (TMM) (Robinson and Oshlack, 2010) using edgeR (v. 3.20.9) (Robinson et al., 2010). Specifically, RUVs (with k = 5) was used after defining groups based on both genotype and treatment and using a list of 2677 negative control genes obtained as genes with an adjusted p-value > 0.9 in the comparison between sleep deprivation and controls in microarray data available through GEO (GSE78215) (obtained from (Gerstner et al., 2016)). Analogously, a list of 579 positive control genes to evaluate the effect of normalization was obtained from the same study.

Differential expression analysis was performed using edgeR (v. 3.20.9) with a factorial design that included genotype (WT or Shank3^∆C^) and treatment (homecage control or sleep deprivation). We specified the following contrasts: wild type sleep deprived vs wild type homecage controls (WTSD vs. WTHC); Shank3^∆C^ sleep deprived vs Shank3^∆C^ homecage controls (S3SD vs. S3HC); Shank3^∆C^ controls vs wild type controls (S3HC vs. WTHC); and Shank3^∆C^ sleep deprived vs wild type sleep deprived (S3HC vs. WTHC). To gain insight on the different effects of sleep deprivation between wild type and mutant, we considered the union of the differentially expressed genes between genotypes both in controls and sleep deprived mice. We clustered the genes using k-means (k = 3) and the samples using hierarchical clustering based on the selected genes. The resulting ordered genes and samples were visualized as a heatmap using the pheatmap package (v. 1.0.10). The R code used for RNA-seq data analysis is publicly available at https://github.com/drighelli/peixoto.

Functional annotation analysis of genes differentially expressed at FDR < 0.1 was based on Ensembl Gene IDs and performed using the database for annotation, visualization, and integrated discovery v 6.7 (DAVID, https://david.ncifcrf.gov). The following functional categories were used: GO Biological Process, GO Molecular Function, Uniprot keywords and KEGG pathways. Enrichment was determined relative to all genes expressed in the mouse prefrontal cortex based on our data. We used an enrichment p-value (EASE) cutoff < 0.1 for individual functional terms. Enriched functional terms were clustered at low stringency, to obtain clusters with an enrichment score > 1.2 (corresponding to an average EASE > 0.05).

### Assessment of circadian rhythms

#### Experimental design

Adult male (9-week-old) wild type (n = 8) and Shank3^∆C^ (n = 7) littermate mice were individually housed with a low-profile wireless running wheel (Med Associates Inc, Fairfax, VT) and kept under 12:12 h light:dark (LD) cycle for 3 weeks (including 1 week of habituation) followed by 3 weeks of complete darkness (DD). Two Shank3^∆C^ were excluded from the experiemnts because they failed run on the wheel for the duration of the experiment.

#### Data acquisition and analysis

Wheel running behavior was continuously monitored using wheel managing software and exported into 10 min time bins (Med Associates Inc). ClockLab version 6.0.50 (Actimetrics, Wilmette, IL) was used for analysis. Period, the circadian length in time from activity onset to the next activity onset, was calculated using ClockLab’s chi-squared periodogram analysis function. Alpha, the length of activity period, was calculated by the time between activity onset and offset (defined by manufacturer defaults). Activity was calculated as the total sum of wheel revolutions per day. Comparisons between genotypes (WT vs. Shank3^∆C^) were done with paired or unpaired Student’s t-test as appropriate for period length, alpha length, and wheel running activity. A linear mixed-effects model was used to estimate the contributions of genotype and the interaction between genotype and time for wheel running activity for LD week 3 through DD week 3. The model was fitted using genotype, time, and their interaction as fixed effects, and the individual mice were labeled as a random effect. The analysis was conducted with the R package nlme (v. 3.5.0) and is publicly available at https://github.com/drighelli/peixoto.

## Supplementary materials (list)

Figure 1 – table supplement 1. Incidence of sleep problems in Autism Spectrum Disorder (ASD) compared to typically developing (TD) individuals.

Figure 2 – figure supplement 1. Baseline sleep bout analysis.

Figure 2 – table supplement 1. EEG spectral analysis statistics.

Figure 3 – figure supplement 1. Shank3^∆C^ mice sleep less after sleep deprivation.

Figure 3 – figure supplement 2. Sleep bout analysis after sleep deprivation.

Figures 2 and 3 – table supplement 1. Source data used for generating plots and statistics for Figures 2 and 3.

Figure 4 – table supplement 1. Genes differentially expressed in Shank3^∆C^ vs. WT mice.

Table 1 – table supplement 1. Functional annotation clustering analysis of genes in Figure 4 – table supplement 1.

Figure 4 – file supplement 1. Detailed report of statistical analysis including R code to reproduce the analysis.

Figure 5 – figure supplement 1. Actograms for all wild type and Shank3^∆C^ mice.

Table 2 – table supplement 1. Results of genotype × time interaction analysis for LD week 3 through DD week 3.

## Article and author information

### Author details

Ashley M Ingiosi. Department of Biomedical Sciences, Elson S. Floyd College of Medicine, Washington State University, Spokane, Washington, United States. **Contribution:** Experimental design, data collection, data analysis, manuscript writing and editing. **Competing interests:** No competing interests declared

Taylor Wintler. Department of Biomedical Sciences, Elson S. Floyd College of Medicine, Washington State University, Spokane, Washington, United States. **Contribution:** Experimental design, data collection, data analysis, manuscript writing and editing. **Competing interests:** No competing interests declared

Hannah Schoch. Department of Biomedical Sciences, Elson S. Floyd College of Medicine, Washington State University, Spokane, Washington, United States. **Contribution:** Data collection, data analysis, manuscript writing and editing. **Competing interests:** No competing interests declared

Kristan G Singletary. Department of Biomedical Sciences, Elson S. Floyd College of Medicine, Washington State University, Spokane, Washington, United States. **Contribution:** Data collection, manuscript editing. **Competing interests:** No competing interests declared

Dario Righelli. Istituto per le Applicazioni del Calcolo “M. Picone”, Consiglio Nazionale della Ricerche, Napoli, Italy. Dipartimento di Scienze Aziendali Management & Innovative Systems, Università di Fisciano, Fisciano, Italy. **Contribution:** Data analysis, statistics, manuscript editing. **Competing interests:** No competing interests declared

Leandro G Roser. Department of Biomedical Sciences, Elson S. Floyd College of Medicine, Washington State University, Spokane, Washington, United States. **Contribution:** Data analysis, statistics, manuscript editing. **Competing interests:** No competing interests declared.

Davide Risso. Department of Statistical Sciences, University of Padova, Padova, Italy and Division of Biostatistics and Epidemiology, Department of Healthcare Policy and Research, Weill Cornell Medicine, New York, New York, United States. **Contribution:** Experimental design, data analysis, statistics, manuscript editing. **Competing interests:** No competing interests declared.

Marcos G Frank. Department of Biomedical Sciences, Elson S. Floyd College of Medicine, Washington State University, Spokane, Washington, United States. **Contribution:** Experimental design, manuscript writing and editing. **Competing interests:** No competing interests declared

Lucia Peixoto. Department of Biomedical Sciences, Elson S. Floyd College of Medicine, Washington State University, Spokane, Washington, United States. **Contribution:** Experimental design, data collection, data analysis, manuscript writing and editing. **Competing interests:** No competing interests declared.

## Acknowledgements

Data used in the preparation of this article were obtained from the Phelan-McDermid Syndrome International Registry (PMSIR) version dated 12/1/2016.

We thank Dr. Paul Worley for providing the Shank3 mutant mouse line. We thank Dr. Hans Van Dongen and Dr. John Hogenesch for valuable discussion of the results. We thank Dr. Claudia Angelini for valuable discussion of the statistical methods.

## Ethics

Animal experimentation: This study was performed in strict accordance with the recommendations of the Guide for the Care and Use of Laboratory Animals of the Nation Institutes of Health. All of the animals were handled according to protocols approved by the Washington State University Institutional Animal Care and Use Committee (#04704 and #04705).

## Data and materials availability

Sequencing data generated in this study is available through GEO (GSE113754, secure token for reviewer: ojqlkusoflijnmj). The R code to reproduce all the figures and tables related to RNA-seq data analysis of the article is available on GitHub (github.com/drighelli/peixoto).

## Supplementary materials

**Figure 1- table supplement 1.**
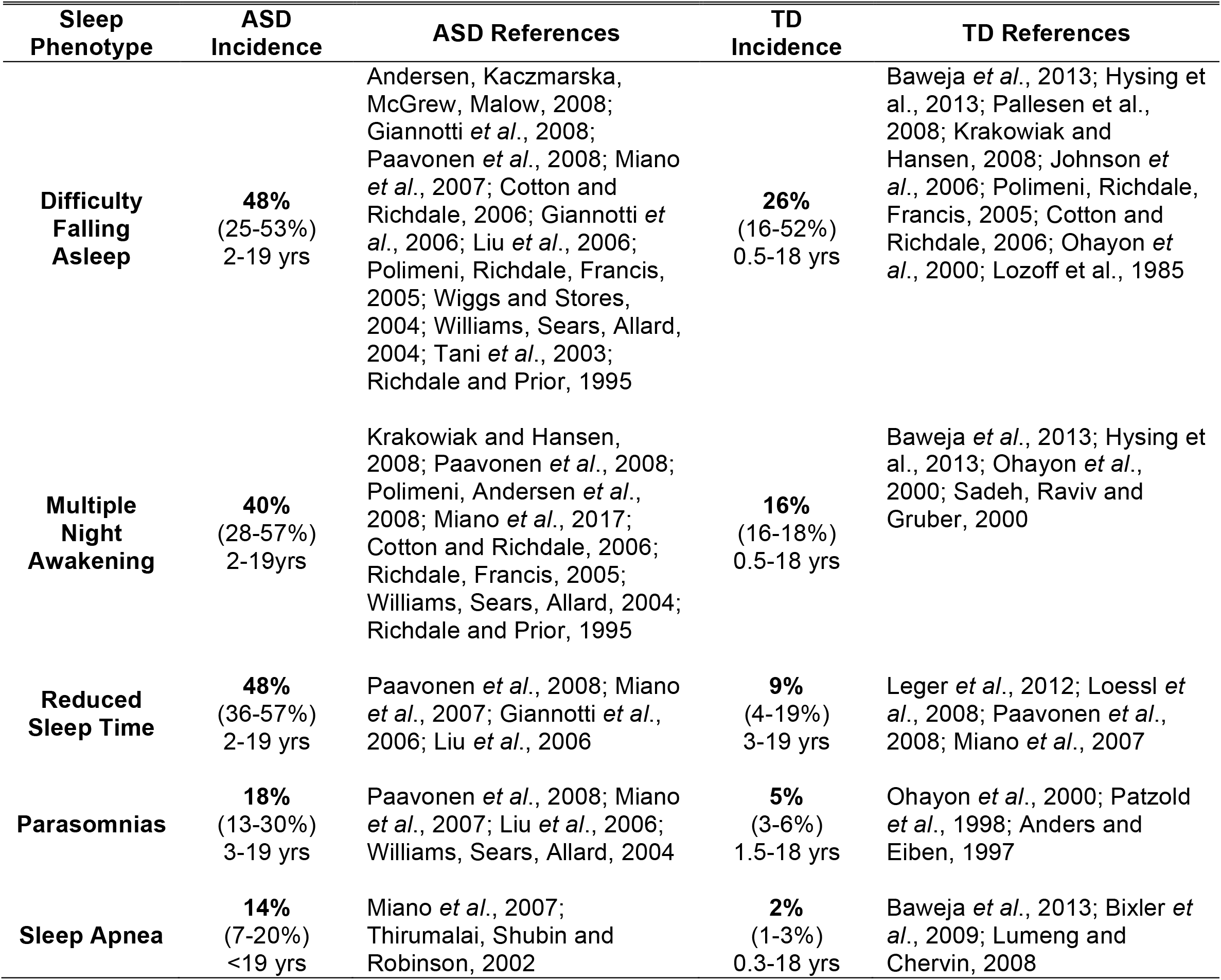
Incidence of sleep problems in Autism Spectrum Disorder (ASD) compared to typically developing (TD) individuals. Meta-analysis of literature, regardless of age. Median, first and third quartile, age range, and included references reported for each phenotype.

**Figure 2- figure supplement 1.**
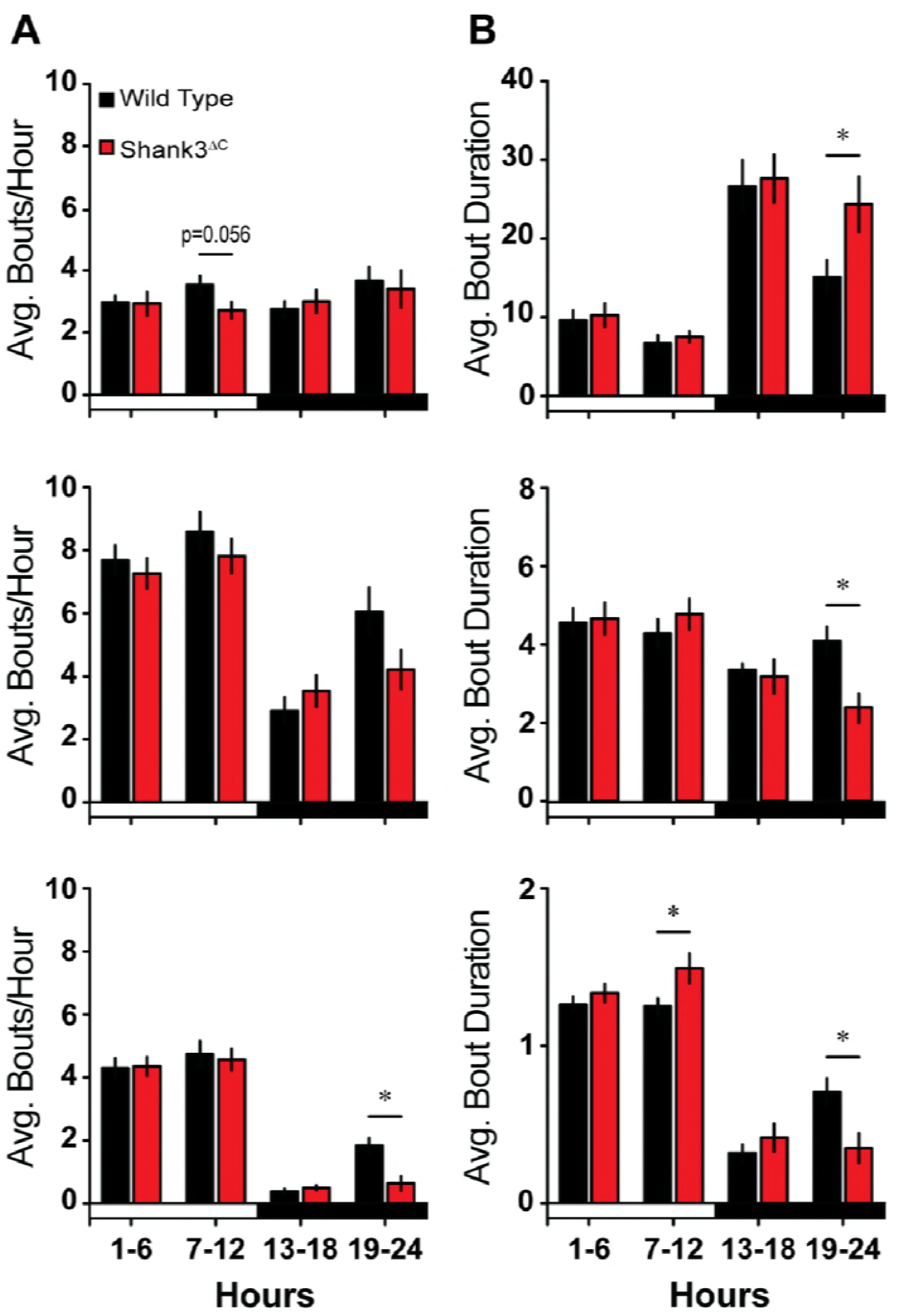
Baseline sleep bout analysis. (**A**) Average number of state-specific bouts per hour (difference scores) shown in 6 h blocks. (**B**) Average bout duration (min) per hour (difference scores) shown in 6 h blocks. Values are means ± SEM for wild type (+/+; n = 10) and Shank3^ΔC^ (-/-; n = 10) mice.

**Figure 2- supplement table 1.**
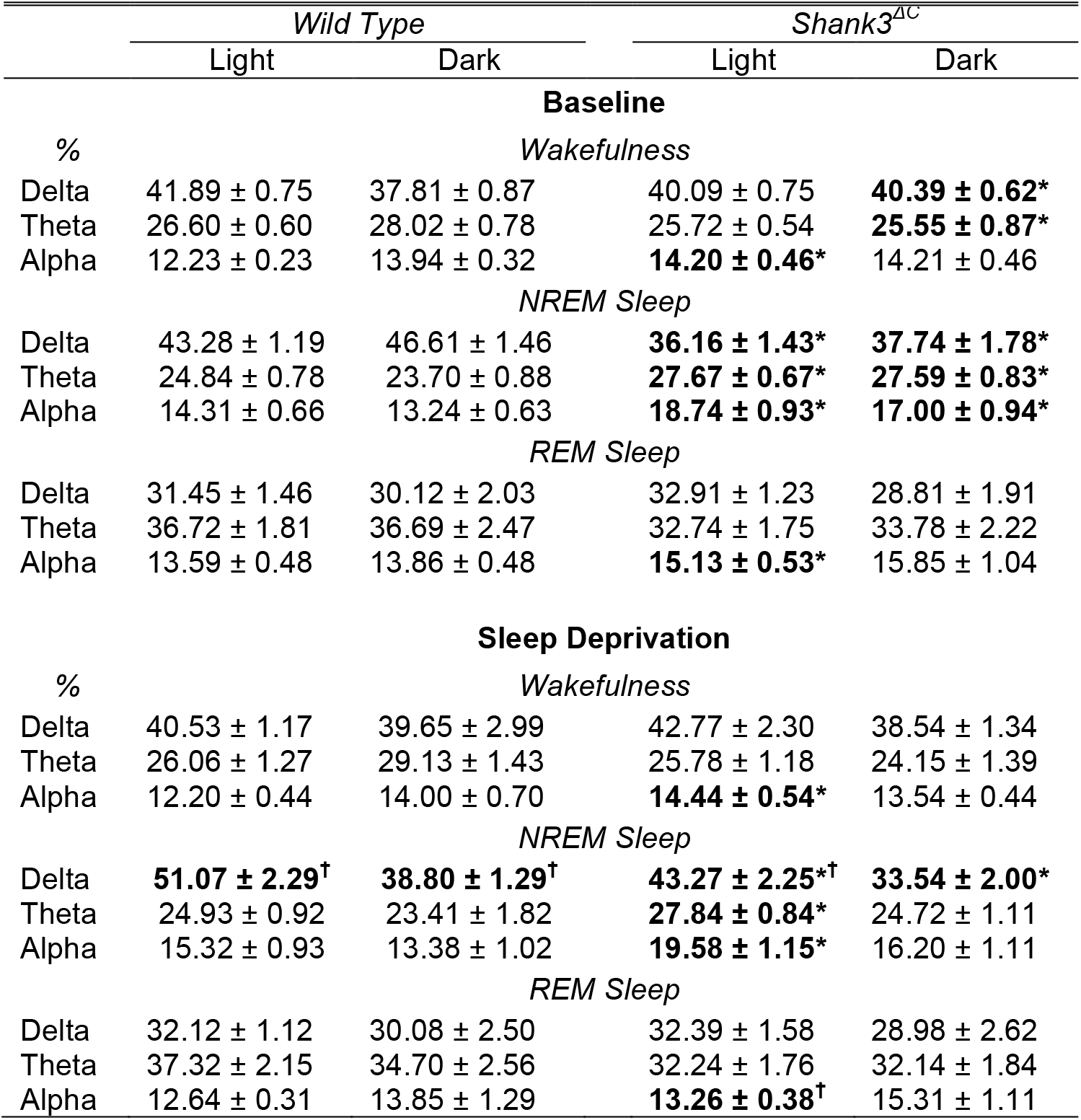
EEG spectral analysis statistics. Shank3^ΔC^ mice exhibit altered EEG spectral characteristics during baseline sleep-wake behavior and after sleep deprivation. Normalized spectral bands for delta (0.5 – 4 Hz), theta (5 – 9 Hz), and alpha (10 – 15 Hz) during wakefulness, NREM sleep, and REM sleep for the light and dark periods expressed as a percentage of the total spectral sum for the light and dark periods, respectively. Analyses between baseline and sleep deprivation light periods for NREM and REM sleep compared baseline hours 1-12 and sleep deprivation hours 6-12. Values are means ± SEM for wild type (n = 10) and Shank3^ΔC^ (n = 10). *p<0.05; difference from respective wild type light and dark period values. ^†^p<0.05; difference from respective baseline light and dark period values.

**Figure 3- figure supplement 1.**
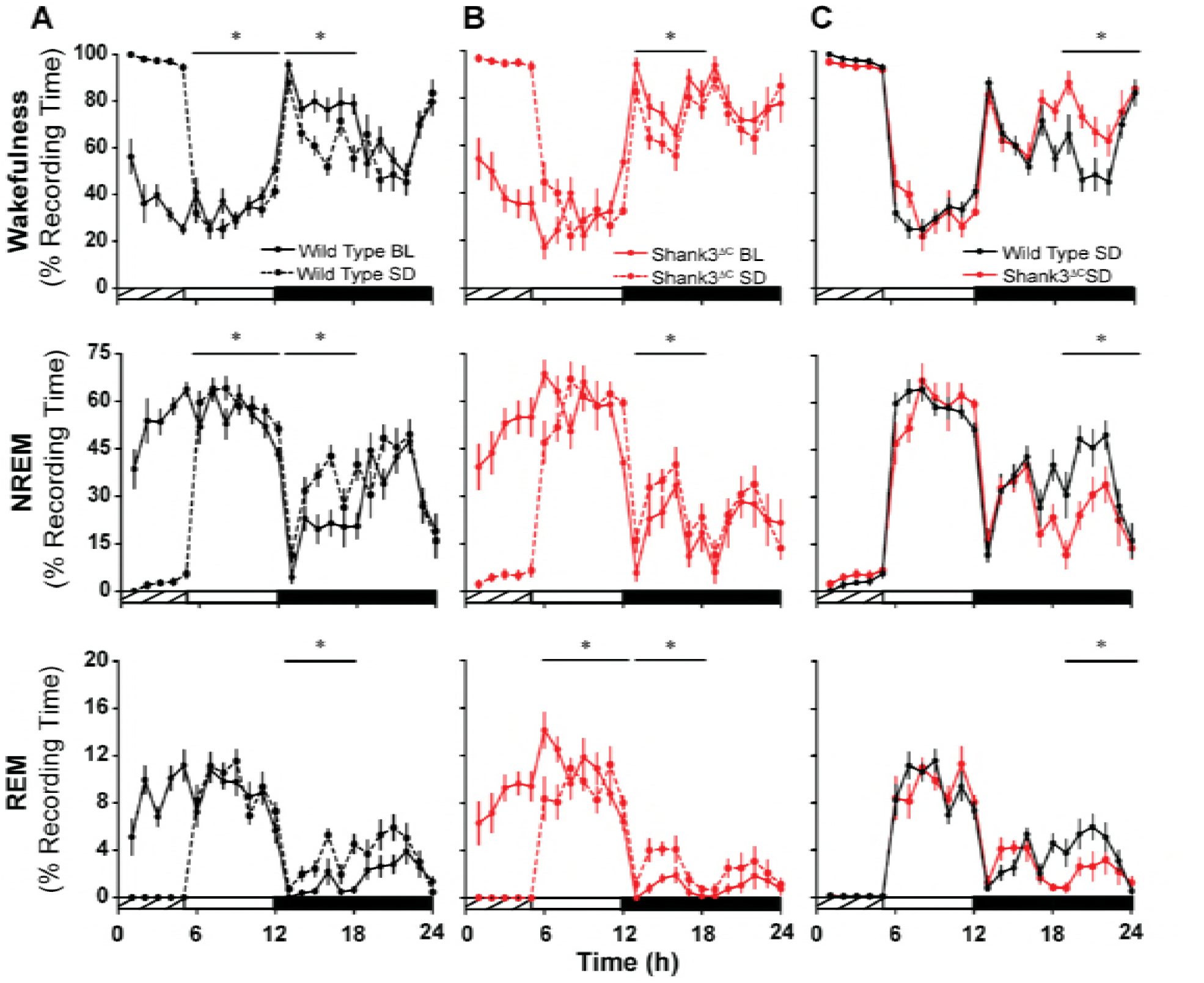
Shank3^ΔC^ mice sleep less after sleep deprivation. The rows represent the arousal states of wakefulness (top), NREM sleep (middle), and REM sleep (bottom). (**A-B**) Time in state shown as percentage of recording time per hour comparing baseline and sleep deprived conditions for wild type mice (**A**) and Shank3^ΔC^ mice (**B**). (**C**) Time in state shown as percentage of recording time per hour under sleep deprived conditions. The cross-hatched bars on the x-axis denotes the 5 h sleep deprivation period, the open bars denotes the light period, and the filled bars denotes the dark period of the light:dark cycle. Values are means ± SEM for wild type (+/+; n = 10) and Shank3^ΔC^ (-/-; n = 10) mice, *p < 0.05.

**Figure 3- figure supplement 2.**
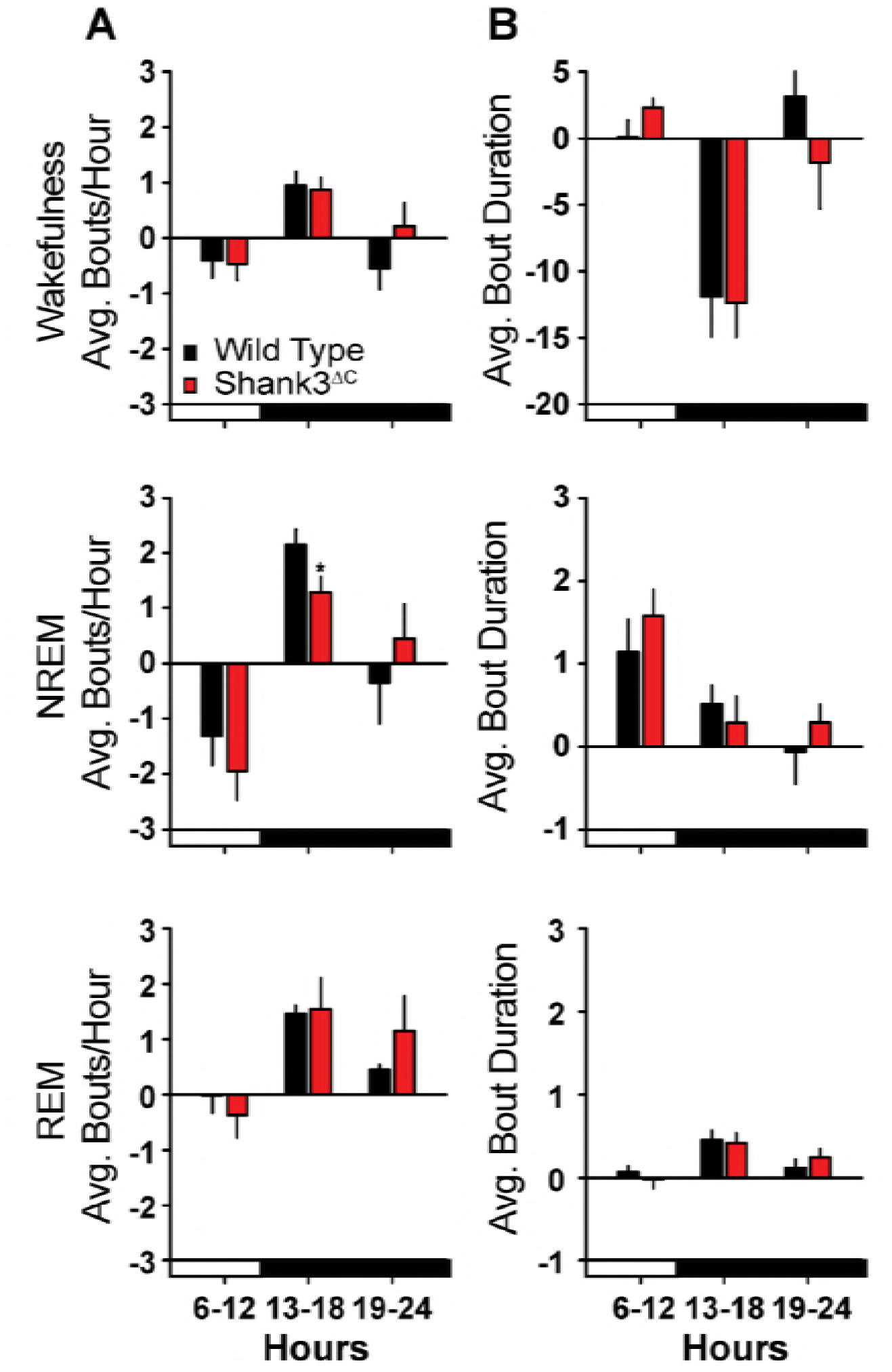
Sleep bout analysis after sleep deprivation. (**A**) Average number of state-specific bouts per hour (difference scores) shown in 7 h (remaining light period) and 6 h blocks. (**B**) Average bout duration (min) per hour (difference scores) shown in 7 h (remaining light period) and 6 h blocks. Difference scores were calculated by subtracting baseline (BL) values from sleep deprivation (SD) values. Values are means ± SEM for wild type (+/+; n = 10) and Shank3^ΔC^ (-/-; n = 10) mice, *p < 0.05.

**Figures 2 and 3 – table supplement 1**. Source data used for generating plots and statistics for Figures 2 and 3.

**Figure 4 - supplement table 1.** Genes differentially expressed between Shank3^∆C^ vs. WT mice. RNA-seq study of gene expression from cortex of obtained from adult male Shank3^ΔC^ and wild type (WT) mice, either under control home cage conditions (HC) or following 5 hours of sleep deprivation (SD). N=5 mice per group. False discovery rate (FDR) <0.1.

**Table 1- supplement table 1.** Functional annotation clustering analysis of genes in Figure 4- supplement table 1. Functional annotation and clustering analysis was performed using DAVID (https://david.ncifcrf.gov) and functional information from the following databases: GO (Biological process and Molecular function), KEGG pathways, Uniprot keywords. Enrichment was performed relative to all transcripts expressed in the mouse pre-frontal cortex as defined by our RNA-seq data. Enriched functional terms where clustered at low stringency, to obtain clusters with enrichment score > 1.2 (corresponding to an average p-value > 0.05).

**Figure 4 – file supplement 1.** Detailed report of statistical analysis including R code to reproduce the analysis.

**Figure 5- supplement figure 1.**
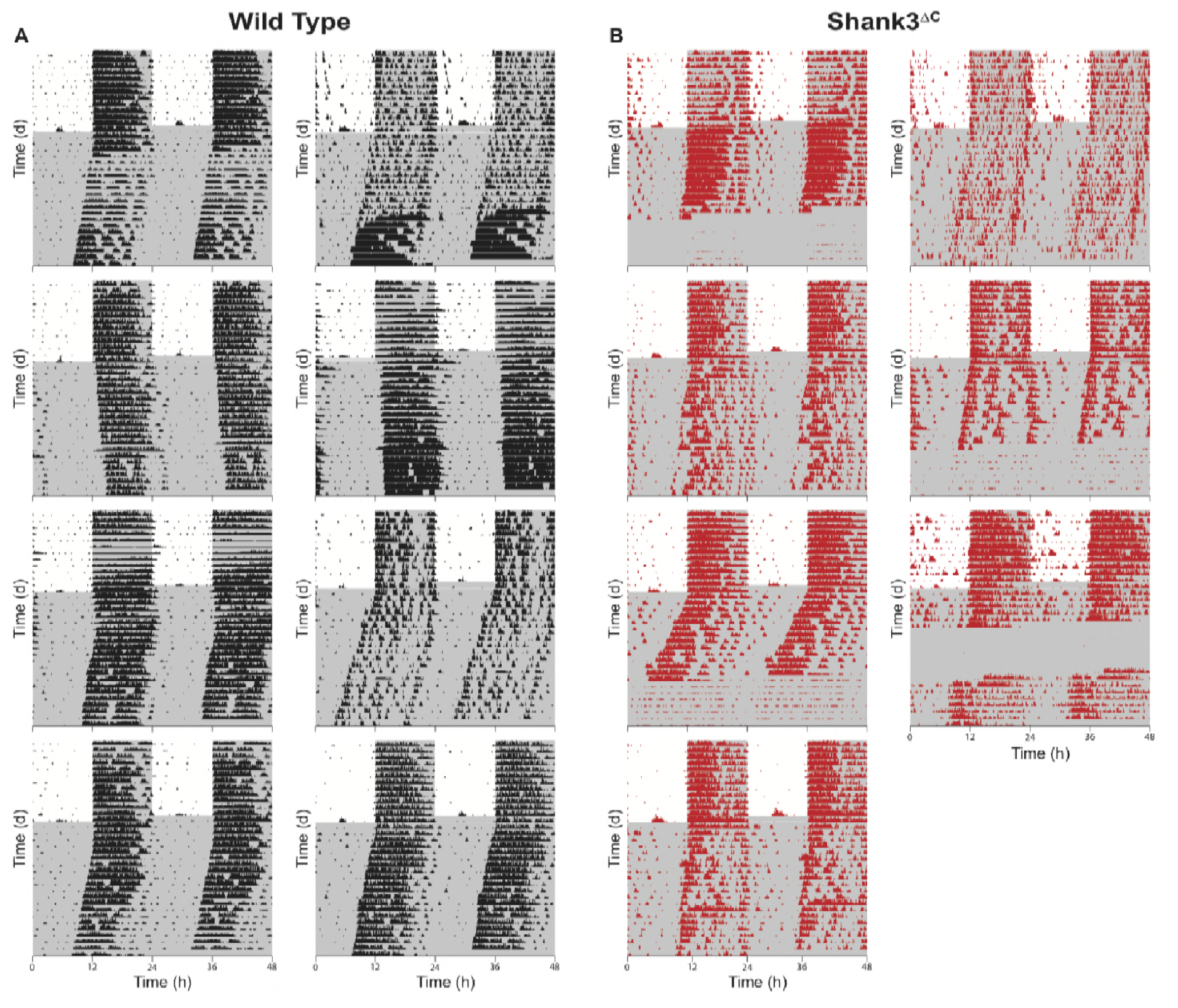
Actograms for all wild type and Shank3^∆C^ mice. Mice were entrained to a 12:12 light dark cycle (LD, 559.0±4.0:0±0.0Lux) for two weeks prior to 3 weeks constant darkness (DD, 0±0.0Lux). Gray shading is representative of the dark period. **(A)** Actograms for wild type mice. **(B)** Actograms for Shank3^∆C^ mice. Wild type (+/+; n = 8) and Shank3^∆C^ (-/-; n = 7) mice.

**Table 2- supplement table 1.** Results of genotype × time interaction analysis for LD week 3 through DD week 3. Linear mixed-effects with interaction genotype × time for period, alpha, and running wheel activity for LD week 3 through DD week 3. The model was fitted using genotype, time, and their interaction as fixed effects, and the individual mice labeled as random effect. There is a significant effect for genotype × time and activity (p<0.0001, ANOVA) and no significant effect for genotype × time or genotype for period and alpha. Significance at p<0.05. numDF, numerator degrees of freedom. denDF, denominator degrees of freedom.

|  | numDF | denDF | F-value | p-value |
| --- | --- | --- | --- | --- |
| **Period** |  |  |  |  |
| (Intercept) | 1 | 39 | 351.35 | **<0.0001** |
| Week | 3 | 39 | 3.36 | **0.03** |
| Genotype | 1 | 13 | 0.01 | 0.94 |
| Genotype x Week | 3 | 39 | 1.90 | 0.15 |
| **Alpha** |  |  |  |  |
| (Intercept) | 1 | 373 | 909.26 | **<0.0001** |
| Time | 1 | 373 | 2.17 | 0.14 |
| Genotype | 1 | 13 | 2.12 | 0.17 |
| Genotype x Time | 1 | 373 | 0.02 | 0.88 |
| **Activity** |  |  |  |  |
| (Intercept) | 1 | 388 | 52.83 | **<0.0001** |
| Time | 1 | 388 | 0.33 | 0.57 |
| Genotype | 1 | 13 | 6.77 | **0.02** |
| Genotype x Time | 1 | 338 | 30.96 | **<0.0001** |

